# Unrecognized Potent Activities of Colistin Against Clinically Important *mcr*+ Enterobacteriaceae Revealed in Synergy with Host Immunity

**DOI:** 10.1101/2023.03.21.533661

**Authors:** Monika Kumaraswamy, Angelica Riestra, Anabel Flores, Satoshi Uchiyama, Samira Dahesh, Gunnar Bondsäter, Victoria Nilsson, Melanie Chang, Hideya Seo, George Sakoulas, Victor Nizet

**Affiliations:** Division of Infectious Diseases and Global Public Health, Department of Medicine, UC San Diego, La Jolla, CA, USA; Infectious Diseases Section, VA San Diego Healthcare System, San Diego, CA, USA; Department of Biology, San Diego State University, San Diego, CA, USA; Division of Host-Microbe Systems and Therapeutics, Department of Pediatrics, UC San Diego, La Jolla, CA, USA; Department of Biological Sciences, California Baptist University, Riverside, CA, USA; Faculty of Medicine, Lund University, Lund, Sweden; School of Medicine, China Medical University, Taichung, Taiwan; Department of Anesthesia, Kyoto University, Kyoto, Japan; Sharp Rees Stealy Medical Group, San Diego, CA, USA; Skaggs School of Pharmacy and Pharmaceutical Sciences, UC San Diego, La Jolla, CA, USA

## Abstract

Colistin (COL) is a cationic cyclic peptide that disrupts negatively-charged bacterial cell membranes and frequently serves as an antibiotic of last resort to combat multidrug-resistant Gram-negative bacterial infections. Emergence of the horizontally transferable plasmid-borne mobilized colistin resistance (*mcr*) determinant and its spread to Gram-negative strains harboring extended-spectrum β-lactamase and carbapenemase resistance genes threatens futility of our chemotherapeutic arsenal. COL is widely regarded to have zero activity against *mcr+* patients based on standard antimicrobial susceptibility testing (AST) performed in enriched bacteriological growth media; consequently, the drug is withheld from patients with *mcr+* infections. However, these standard testing media poorly mimic in vivo physiology and omit host immune factors. Here we report previously unrecognized bactericidal activities of COL against *mcr-1+* isolates of *Escherichia coli* (EC), *Klebsiella pneumoniae* (KP), and *Salmonella enterica* (SE) in standard tissue culture media containing the physiological buffer bicarbonate. Moreover, COL promoted serum complement deposition on the *mcr-1+* Gram-negative bacterial surface and synergized potently with active human serum in pathogen killing. At COL concentrations readily achievable with standard dosing, the peptide antibiotic killed *mcr-1+* EC, KP, and SE in freshly isolated human blood proved effective as monotherapy in a murine model of *mcr-1+* EC bacteremia. Our results suggest that COL, currently ignored as a treatment option based on traditional AST, may in fact benefit patients with *mcr-1+* Gram negative infections based on evaluations performed in a more physiologic context. These concepts warrant careful consideration in the clinical microbiology laboratory and for future clinical investigation of their merits in high risk patients with limited therapeutic options.

## INTRODUCTION

The non-ribosomally synthesized cationic polypeptide antibiotic colistin (COL, polymyxin E) was first isolated from the soil bacterium *Paenibacillus polymyxa* subsp. *colistinus* in 1949 (1). COL has dose-dependent bactericidal activity against most Gram-negative bacteria, but its usage was largely abandoned in the 1970s due to associated nephro- and neurotoxicity and the emergence of less toxic antibiotic alternatives such as cephalosporins, aminoglycosides, and quinolones. However, an increasing prevalence of multidrug-resistant (MDR) Gram-negative infections caused by carbapenem-resistant *Enterobacteriaceae* (CRE), *Acinetobacter baumannii* (AB) and *Pseudomonas aeruginosa* (PA) with limited therapeutic options has prompted a resurgence of COL usage as a last line of defense (2, 3) and its designation by the World Health Organization as a critically important antimicrobial for human medicine (4). As COL resistance determinants emerge in CRE and other highly MDR Gram-negative bacteria (5–8), a prospect of strains resistant to all conventional antibiotics is widely feared.

A multicomponent polypeptide comprised of a hydrophilic cyclic heptapeptide, tripeptide side chain, and a hydrophobic acylated fatty acid residue (6-methyl-octanoic acid or 6-methyl-heptanoic acid), COL binds to lipopolysaccharides (LPS) on the outer membrane (OM) of target Gram-negative bacteria (9). Like other polymyxins, COL contains cationic α,γ-di-aminobutyric acid residues that competitively displace LPS-stabilizing divalent cations (Mg^+2^ and Ca^+2^) from negatively-charged phosphate groups, leading to cell envelope permeabilization, leakage of intracellular contents, and bacterial cell death (9). The primary mechanisms of bacterial resistance to COL involve covalent cationic modifications to LPS structure, including the addition of phosphoethanolamine, 4-amino-4-deoxy-L-arabinose, and/or galactosamine to the phosphate group of lipid A or core oligosaccharide (10). These alterations neutralize LPS negative charge thereby diminishing COL binding affinity to its target (9).

Historically, genetic determinants of COL resistance were localized to the chromosome, impeding widespread dissemination through and across bacterial populations and limiting clinical impact (10). This changed dramatically upon the emergence of the horizontally transferable plasmid-mediated mobilized colistin resistance (*mcr-1*) determinant encoding an LPS-modifying transferase enzyme that adds phosphoethanolamine to lipid A. First identified in 2015 among *Escherichia coli* strains in pigs and humans in China (11), *mcr-1* has subsequently spread rapidly among environmental, animal and human clinical isolates worldwide to pose a severe global public health threat (12, 13). Furthermore, numerous studies have documented diversification of a large array of plasmid-borne *mcr* variants, from *mcr-2* to *mcr-10* (14–20), highlighting the success of this resistance paradigm to evolutionary selective forces. Spread of *mcr-1* and variants now encompasses Gram-negative bacteria harboring extended-spectrum ®- lactamase and carbapenemase resistance genes, threatening the therapeutic utility of our full existing antibiotic armamentarium (5–8).

COL is never recommended in the therapy of Gram-negative bacteria upon molecular detection of *mcr-1* or related plasmids. This medical decision making reflects consistent elevated minimum inhibitory concentration (MIC) values identified by antimicrobial susceptibility testing (AST) in the clinical microbiology laboratory. AST is routinely performed using a nutrient-rich bacteriological media, specifically cation-adjusted Mueller-Hinton broth (CA-MHB), using standardized protocols and breakpoints developed and regularly updated by the Clinical and Laboratory Standards Institute (CLSI) in the United States (21) and the European Committee on Antimicrobial Susceptibility Testing (EUCAST) (22). However, when faced with MDR bacterial isolates exhibiting few if any drug susceptibilities and a high incidence of clinical treatment failure, we (23–25) and others (26–28) have advocated greater circumspection regarding key limitations of allowing a single “gold standard” *in vitro* AST assay to guide antibiotic selection. Firstly, AST is performed in media (CA-MHB) composed of beef extract, casein, and starch, chosen for optimal bacterial growth—a molecular composition entirely distinct from that of infected human tissues and fluids where the bacterial targets of antibacterial action are located. Secondly, AST incorporates no molecular or cellular elements of host innate immunity such as endogenous antimicrobial peptides (AMPs), serum complement, or phagocytes, which may interact with any given pharmacological antibiotic synergistically or antagonistically. Recent studies have shown that diverse drugs including ampicillin (29), nafcillin (30), azithromycin (24, 31), rifabutin (32), and avibactam (33), to which MDR strains of key human bacterial pathogens are deemed highly resistant by standard MIC testing, do indeed exhibit potent (but neglected) antimicrobial activities against these same strains when AST is performed in media that closely mimics human physiology or in synergy with host immune factors.

Given the urgent concern of plasmid-borne COL resistance now spreading within the highest threat MDR Gram-negative bacterial pathogens, we chose to investigate the impact of more physiological media conditions and host defense components on the susceptibility of various medically important Enterobacteriaceae harboring *mcr* gene variants (*mcr-1* to *mcr-4*) to COL and the related cationic peptide antibiotic polymyxin B (PMB). Our *in vitro, ex vivo*, and *in vivo* studies in this context uncover clear, and likely therapeutically meaningful, COL and PMB activities against diverse *mcr*+ Gram-negative bacterial pathogens. These activities are currently hidden from practitioners who remain reliant on standard AST to guide management of critically ill patients, a matter we propose deserves careful attention and further clinical investigation.

## RESULTS

### Polymyxins retain activity against mcr+ Gram-negative bacteria in more physiological media testing conditions

We compared the MIC of COL and PMB against 12 strains of Gramnegative bacteria (7 *E. coli*, 4 *Salmonella* spp., 1 *Klebsiella pneumoniae*), each harboring plasmid-mediated colistin-resistance (*mcr-1* to *mcr-4*) genes in (a) standard bacteriological testing medium (CA-MHB) versus (b) a more physiological medium based on the common mammalian tissue culture medium Roswell Park Memorial Institute (RPMI) 1640 (34). To ensure equivalent bacterial growth kinetics, the RPMI 1640 medium was supplemented with 10% Luria broth and designated RPMI(10%LB) (24). A strong multifold reduction in COL and PMB MICs were observed for all isolates tested in RPMI(10%LB) compared to CA-MHB, with the COL MIC dropping to ≤ 2 mg/L for 12/12 strains and the PMB MIC dropping to ≤ 2 mg/L for 11/12 strains in the more physiological media (**Table 1)**. Of note, the current clinical MIC breakpoint of COL established by EUCAST for Enterobacterales is ≤ 2 mg/L (22), a concentration readily achievable in serum by standard therapeutic dosing of intravenous colistimethate sodium (35).

**Table 1.** Comparative MIC testing for colistin and polymyxin B performed in standard bacteriological (CA-MHB) and supplemented mammalian tissue culture RPMI(10%LB) media.

| AR Bank Isolate | Bacteria | Resistance Genotype | Colistin (mg/L) |  | Polymyxin B (mg/L) |  |
| --- | --- | --- | --- | --- | --- | --- |
|  |  |  | CA-MHB | RPMI (10% LB) | CA-MHB | RPMI (10% LB) |
| #0346 | *Escherichia coli* | *mcr-1* | 8 | 1 | 4 | 1 |
| #0349 | *Escherichia coli* | *mcr-1* | 4 | 1 | 2 | 1 |
| #0350 | *Escherichia coli* | *mcr-1* | 4 | 1 | 4 | 1 |
| #0493 | *Escherichia coli* | *mcr-1* | 8 | 1 | 8 | 1 |
| #0494 | *Escherichia coli* | *mcr-1* | 8 | 1 | 4 | 4 |
| #0495 | *Escherichia coli* | *mcr-1* | 8 | 1 | 4 | 2 |
| #0496 | *Salmonella* Enteritidis | *mcr-1* | 16 | 1 | 8 | 1 |
| #0497 | *Klebsiella pneumoniae* | *mcr-1* | 16 | 2 | 8 | 2 |
| #0538 | *Escherichia coli* | *mcr-2* | 8 | 1 | 8 | 1 |
| #0539 | *Salmonella* Typhimurium | *mcr-3* | 8 | 1 | 4 | 1 |
| #0540 | *Salmonella* Oslo | *mcr-3* | 8 | 1 | 4 | 1 |
| #0635 | *Salmonella* Typhimurium | *mcr-4* | 16 | 1 | 8 | 1 |

Note: MIC results for each antimicrobial agent for an isolate may commonly be $\pm 1$ log<sub>2</sub> (doubling dilution) different than what is posted on the FDA & CDC AR Bank website because this is the normal technical variability of antimicrobial susceptibility testing (see J. H. Jorgensen. 1993. J Clin Microbiol. Vol 31[11]: 2841-2844).

### COL activity against mcr-1 + Gram-negative bacteria in physiological media is bactericidal, dependent on bicarbonate, and involves membrane permeabilization

Media-dependent COL susceptibility was further examined in kinetic killing assays performed with *mcr-1*-harboring strains of *E. coli* (EC), *K. pneumoniae* (KP) and *S. enterica* (SE), each at an initial inoculum of 5 x 10^5^ CFU/ml. COL at 1 or 2 mg/L showed potent bactericidal activity against all three bacterial strains in the more physiological RPMI(10%LB) media, with no recovered bacteria after 8 h (**Figure 1A**). In stark contrast, the same assay performed in standard bacteriological media CA-MHB found all three *mcr-1+* Gram-negative bacteria achieved rapid logarithmic growth despite the addition of 1 or 2 mg/L COL (**Figure 1A**). Bicarbonate (HCO_3_^-^), the major buffering anion for maintenance of mammalian physiological pH, is a key constituent of RPMI 1640 and other tissue culture media that is lacking in CA-MHB. The presence of HCO_3_^-^ can influence MIC results, sometimes increasing antibiotic potency, e.g. for macrolides or aminoglycosides against several Gram-negative bacteria (24, 36), and other times decreasing antibiotic potency, e.g. for tetracyclines against similar bacterial species (36). We repeated our kinetic killing studies of the *mcr-1+* EC, KP, and SE strains in bicarbonate-free RPMI(10%LB) into which we titrated NaHCO_3_- to achieve lower (10 mM) and higher (25 mM) final concentrations spanning a range characteristic of *in vivo* conditions. Whereas complete bactericidal activity of 1 or 2 mg/L COL against all three species of *mcr-1+* bacteria was seen in the presence of low or high HCO_3_^-^, activity was completely lost in the absence of the anion (**Figure 1B**). Although essential for sensitizing *mcr-1+* bacteria to COL killing in physiological media, HCO_3_^-^ was insufficient to sensitize the same bacteria to COL in the CA-MHB bacteriologic media (**Figure 1B**).

**Figure 1.**
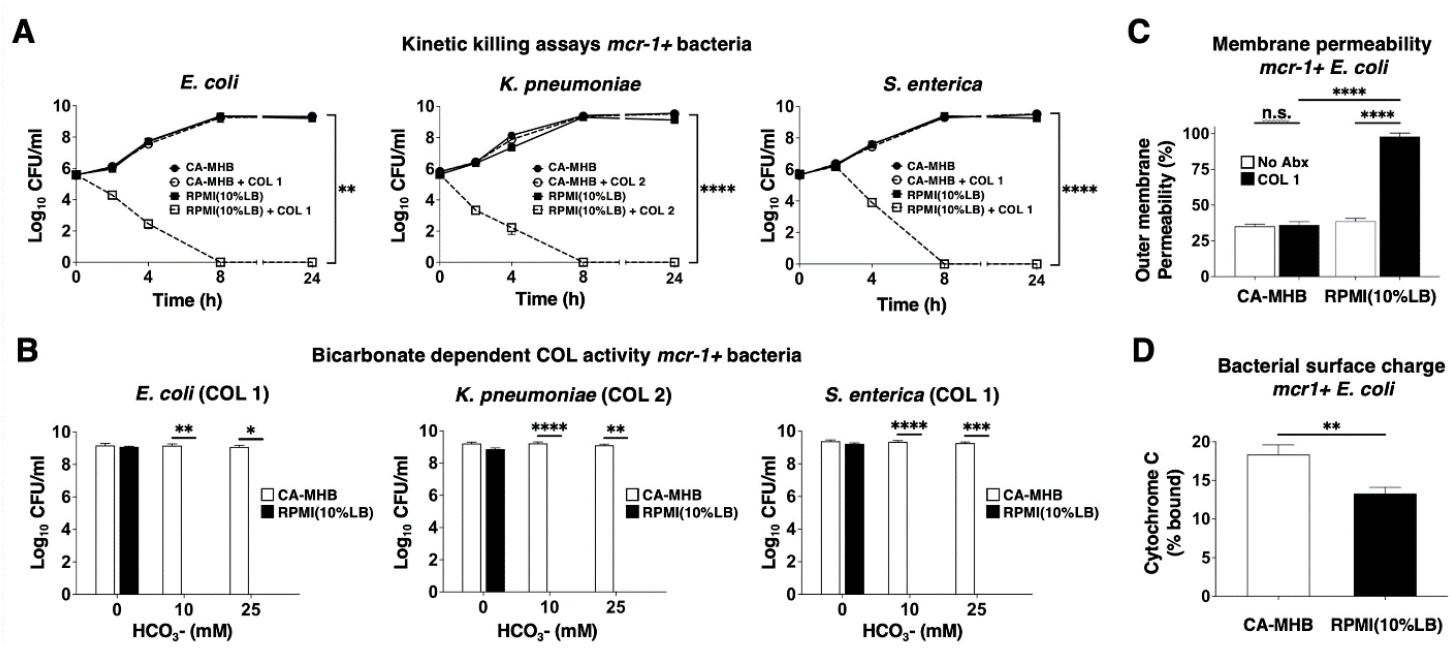
COL bactericidal activity against *mcr-1+* Gram-negative pathogens in physiological media is dependent on bicarbonate and involves membrane permeabilization. (A) Kinetic kill curves show COL minimum bactericidal concentration against *E. coli, K. pneumoniae*, and *S. enterica* (defined as a reduction in viable bacteria ≥3 log_10_ CFU/mL at 24 h vs. starting inoculum) when assessed in supplemented mammalian tissue culture RPMI(10%LB) vs. standard bacteriological medium CA-MHB. (B) Bactericidal activity of COL in RPMI(10%LB) or CA-MHB media amended with 0, 10 or 25 mM HCO_3_-. (C) COL-mediated outer membrane permeabilization of *mcr-1+* EC assessed using the nonpolar compound 1-N-phenylnaphthylamine (NPN) that fluoresces strongly in phospholipid environments in RPMI(10%LB) or CA-MHB. (D) Relative bacterial surface charge estimated by cationic cytochrome c binding to *mcr-1 +* EC following growth in CA-MHB or RPMI(10%LB). Data represent the mean ± SEM from the combination of three experiments performed in triplicate. **P ≤ 0.01 and ****P ≤ 0.0001 by two-way ANOVA.

Changes in EC outer membrane permeabilization upon COL exposure were estimated using the small hydrophobic molecule 1-N-phenylnaphthylamine (NPN) that fluoresces weakly in aqueous environments but strongly when membrane integrity is compromised, binding to bacterial membrane phospholipids after entering the periplasmic space (37). COL treatment of *mcr-1+* EC markedly increased NPN fluorescence (outer membrane permeabilization) in RPMI (10% LB), consistent with its observed bactericidal activity in this media, but produced no distinguishable change from baseline fluorescence when tested in CA-MHB (**Figure 1C**). Increased susceptibility of *mcr-1+* EC to COL in RPMI(10%LB) compared to CA-MHB was not attributable to increased net negative surface charge, as the bacterium bound slightly less cationic cytochrome C in the RPMI(10%LB) than CA-MHB (13% vs. 18%, **Figure 1D**).

### COL accelerates killing of mcr-1+ Gram-negative bacteria in human blood and serum

We next explored if the activity of COL against leading *mcr-1+* Gram-negative bacterial pathogens uncovered in the physiological RPMI(10%LB) media held true in more complex infection-relevant matrices of human blood. Whole blood was freshly collected from normal human volunteers, inoculated with *mcr-1+* strains of EC, KP or SE, with COL added at 0.25x, 0.5x, and 1x the respective MIC identified in RMPI(10% LB), and bacterial CFU enumerated at 1 or 2 h for comparison to the initial inoculum. A significant enhancement of killing of both *mcr-1+* EC and KP was observed upon addition of COL, even at the lowest 0.25x MIC concentration at 1 h (**Figure 2A**). While the *mcr-1+* SE proliferated in human whole blood in the absence of antibiotic, addition of even 0.25x MIC COL significantly reduced recovered CFU over a 2 h time period (**Figure 2A**). A critical element of innate defense against Gram-negative bacteria in blood is the lytic action of serum complement, and recent work indicates that complement can serve to sensitize these bacteria to antibiotics including azithromycin, nisin, avibactam, and vancomycin that are normally considered ineffective based solely on standard AST in bacteriological media (24, 33, 38, 39). Paralleling our findings in whole blood, we identified clear and significant synergy of COL, added as low as 0.25x MIC, to potentiate killing of *mcr-1+* EC, KP, and SE in 10% human serum (**Figure 2B**). Synergy of COL with human serum was extended to the larger panel of 12 Gram-negative bacteria harboring *mcr* plasmids by adding 10% serum to a modified checkerboard MIC assay performed in standard CA-MHB bacteriologic media. For 10 of these isolates, a significantly reduced COL MIC was calculated in the presence of 10% serum **(Table 2**); two strains could not be assessed as they failed to grow in the 10% serum alone. For all strains, 10% heat-inactivated serum did not lower the COL MIC, showing functional complement was required. The MIC of COL dropped ≤ 2 mg/L, the published EUCAST clinical breakpoint for colistin for Enterobacterales, when 10% human serum was added to the standard testing medium.

**Figure 2.**
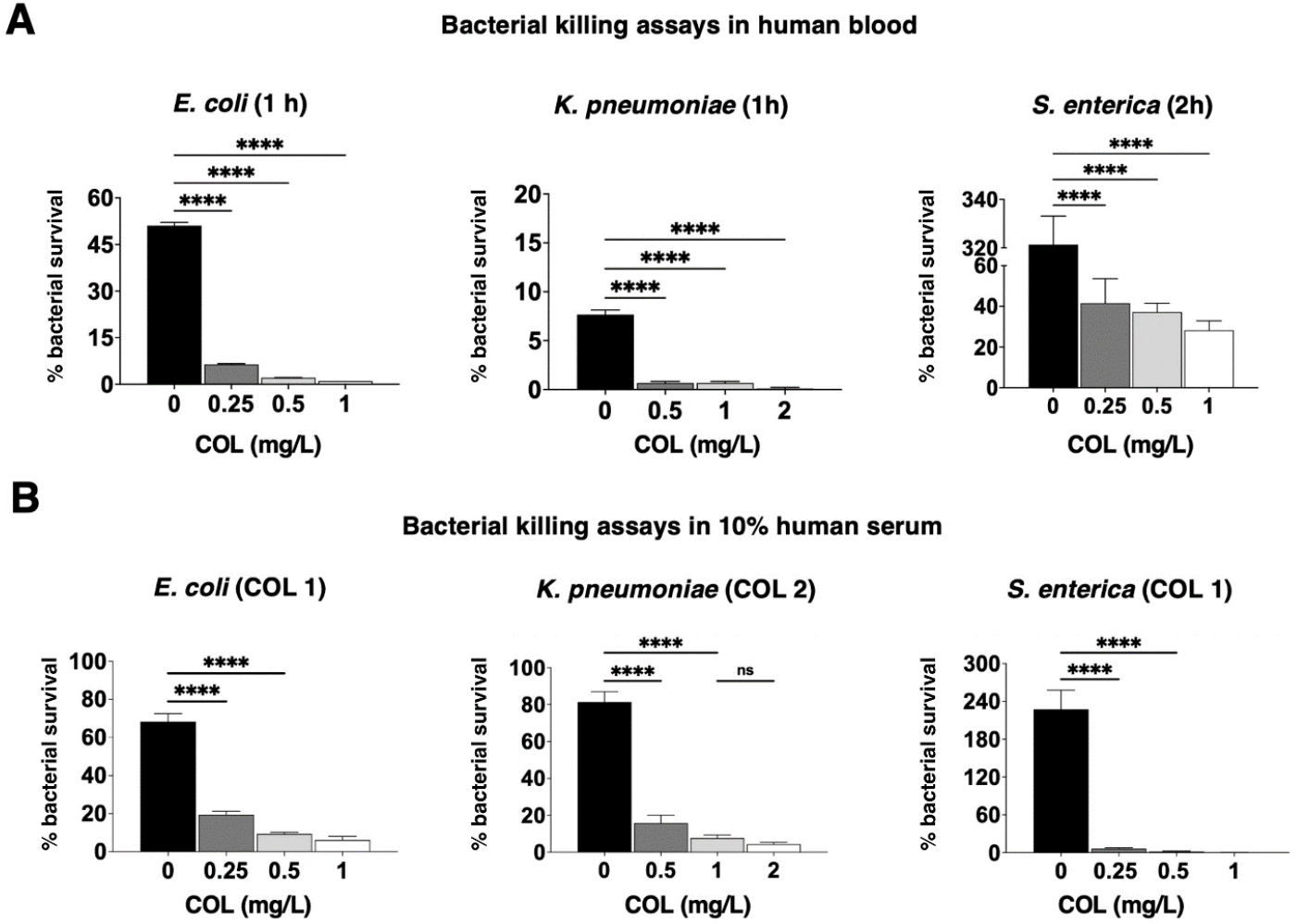
COL accelerates killing of *mcr*-1 + Gram-negative bacteria in human blood and serum. (A) Bacterial survival in freshly isolated human whole blood +/- COL at concentrations representing 1/4x, 1/2x, and 1x MIC for each *mcr-1+* bacterial strain. (B) Bacterial survival in 10% human serum +/- COL/- COL at concentrations representing 1/4x, 1/2x, and 1x MIC for the each *mcr-1+* bacterial strains. Data are represented as % viable CFU vs. initial inoculum and mean ± SEM from a combination of three experiments performed in triplicate as performed using blood or serum from three different donors. ****P ≤ 0.0001 or ns, no statistical significance by one-way ANOVA.

**Table 2.**
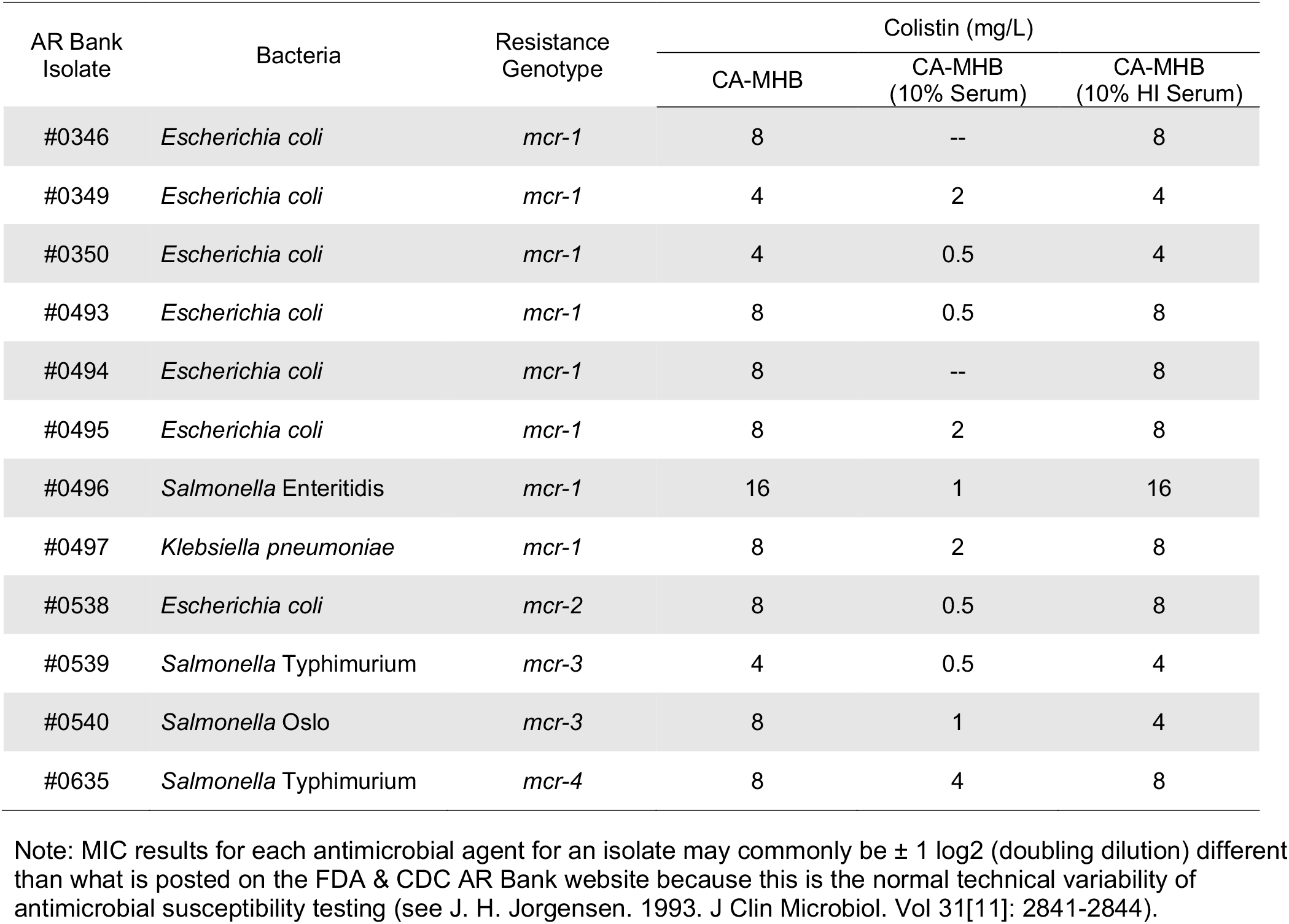
Comparative MIC testing for COL performed in standard bacteriological (CA-MHB) media with or without 10% human serum supplementation.

| AR Bank Isolate | Bacteria | Resistance Genotype | Colistin (mg/L) |  |  |
| --- | --- | --- | --- | --- | --- |
|  |  |  | CA-MHB | CA-MHB (10% Serum) | CA-MHB (10% HI Serum) |
| #0346 | *Escherichia coli* | *mcr-1* | 8 | -- | 8 |
| #0349 | *Escherichia coli* | *mcr-1* | 4 | 2 | 4 |
| #0350 | *Escherichia coli* | *mcr-1* | 4 | 0.5 | 4 |
| #0493 | *Escherichia coli* | *mcr-1* | 8 | 0.5 | 8 |
| #0494 | *Escherichia coli* | *mcr-1* | 8 | -- | 8 |
| #0495 | *Escherichia coli* | *mcr-1* | 8 | 2 | 8 |
| #0496 | *Salmonella* Enteritidis | *mcr-1* | 16 | 1 | 16 |
| #0497 | *Klebsiella pneumoniae* | *mcr-1* | 8 | 2 | 8 |
| #0538 | *Escherichia coli* | *mcr-2* | 8 | 0.5 | 8 |
| #0539 | *Salmonella* Typhimurium | *mcr-3* | 4 | 0.5 | 4 |
| #0540 | *Salmonella* Oslo | *mcr-3* | 8 | 1 | 4 |
| #0635 | *Salmonella* Typhimurium | *mcr-4* | 8 | 4 | 8 |

Note: MIC results for each antimicrobial agent for an isolate may commonly be $\pm 1$ log<sub>2</sub> (doubling dilution) different than what is posted on the FDA & CDC AR Bank website because this is the normal technical variability of antimicrobial susceptibility testing (see J. H. Jorgensen. 1993. J Clin Microbiol. Vol 31[11]: 2841-2844).

### COL promotes C3 deposition on the mcr-1+ Gram-negative bacterial surface

To understand the synergistic killing by COL and serum of *mcr-1+* Gram-negative bacteria, we hypothesized sub-bactericidal COL could promote increased complement deposition and activation on the bacterial surface. All three major pathways of complement activation—the classical, lectin, and alternative pathways, converge at the point of C3 activation (40, 41) and the deposition and activation of C3 on the bacterial cell surface. The bound C3b can be recognized by cognate complement receptors on neutrophils and macrophages for phagocytic uptake (42), or initiate downstream complement cascades culminating in formation of the lytic membrane attack complex (MAC) (43). We assessed C3 binding to the bacterial cell surface using a C3/C3b/C3c antibody by flow cytometry and immunofluorescence microscopy in the presence and absence of COL. At 1x MIC (1 or 2 mg/L) in 20% human serum, COL significantly increased C3 cell surface deposition approximately ≥2-fold on *mcr-1+* EC, KP, and SE compared to bacteria in 20% human serum alone (**Figure 3A**). Quantitative microscopy performed specifically in 10% human serum and 1x MIC of COL for *mcr-1+* EC yielded a 3.5-fold increase in C3 fluorescence signal on the bacterial surface (**Figures 3B and 3C**). Flow cytometry and quantitative microscopy differences between COL-treated and -untreated EC were not seen in control assays performed in 10% heat-inactivated human serum.

**Figure 3.**
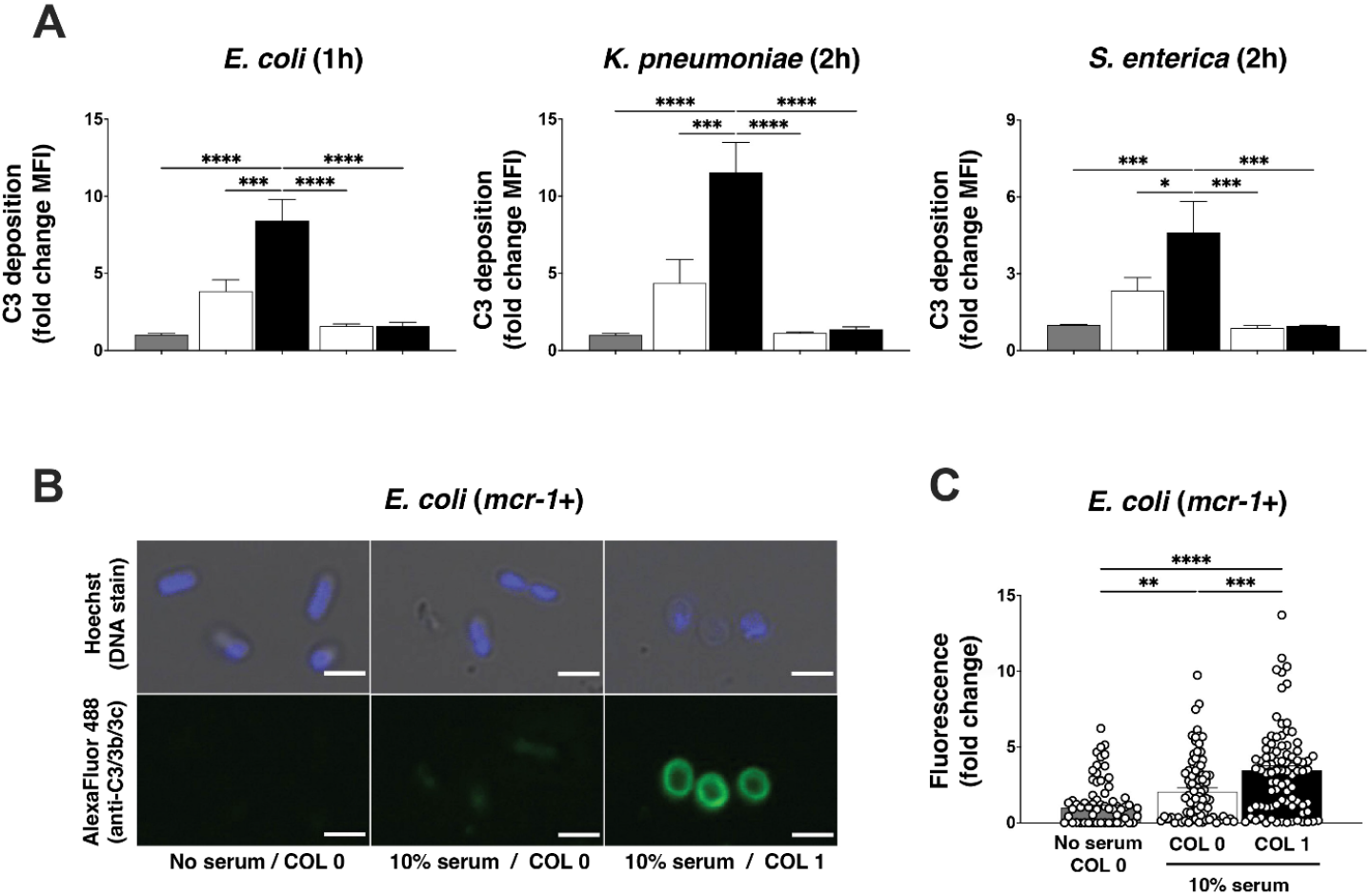
COL promotes C3 deposition on the *mcr-1+* Gram-negative bacterial surface. (A) C3 protein deposition on the surface of E. coli, K. pneumoniae and S. enterica as detected by flow cytometry. Median fluorescence intensity (MFI) of bacteria-bound C3 protein shown as fold change vs. untreated control bacteria of each species. Flow cytometry data representative of two independent experiments conducted in triplicate; 10,000 cells counted per experimental replicate and performed using serum from two different donors. (C) Confocal microscopy images (average intensity Z-projections) of EC in the presence or absence of serum and presence of absence of COL 1 μg/mL. DNA staining by Hoescht dye (blue); bound C3 protein detected using an anti-C3 antibody and fluorescent secondary antibody (Alexa Fluor 488, green) (D) Quantification of C3 fluorescence signal from confocal microscopy shown as fold change vs. untreated control bacteria. Bar graph generated from unbiased analysis of multiple random microscopy fields with >100 cells counted per condition. **P ≤ 0.01, ***P ≤ 0.001 and ****P ≤ 0.0001 by two-way ANOVA.

### COL monotherapy reduces bacterial load and increases survival in a murine model of mcr-1+ E. coli bacteremia

AST performed in standard CA-MHB bacteriological media identifies *mcr-1+* bacteria as resistant to the action of COL, while comparable testing in the more physiological tissue culture-type media RPMI(10% LB) or in the presence of human blood or serum reveals significant “hidden” bactericidal potential of the antibiotic against these same bacteria. To ascertain the *in vivo* relevance of these findings, we intravenously infected C57Bl/6J mice with 1 x 10^9^ CFU of a *mcr-1+* EC strain identified as resistant to COL in CA-MHB but sensitive to COL in RPMI(10% LB), blood or serum. Groups of mice were then treated 1 h post-infection with either a single dose of COL (20 mg/kg) or PBS negative control. As a positive control, a third group of mice received a single 50 mg/kg dose of ceftriaxone (CTX), a commonly prescribed third-generation cephalosporin antibiotic to which this EC strain is sensitive in standard CA-MHB AST. Whereas 85% of the PBS control mice died of infection in the day 10 observation period, mice receiving COL (69% survival) or CTX (61% survival) experienced significant protection from mortality (**Fig. 4A**). To assess bacterial clearance in a nonlethal challenge model, bacteria were treated with 2 x 10^7^ CFU of the *mcr-1+* EC strain then administered the same s.c. antibiotic treatments (COL, CTX, or PBS) at 1 h with a second dose 12 h later. Bacterial CFU recovered from the spleen of mice at 24 h showed a ~15.5-fold reduction with COL treatment and ~12.8-fold reduction with CTX treatment compared to PBS control (**Fig. 4B**). Simultaneous examination of bacterial burden in the kidneys found significant reductions with both antibiotics, with 4 of 6 mice treated with COL and 2 of 6 mice treated with CTX below the detection limit for CFU recovery (**Fig. 4C**). Thus despite the *mcr-1+* EC exhibiting COL resistance by standard MIC testing in CA-MHB, the cationic peptide antibiotic performed equivalently well to the standard of care cephalosporin CTX for treatment of systemic EC infection *in vivo*.

**Figure 4.**
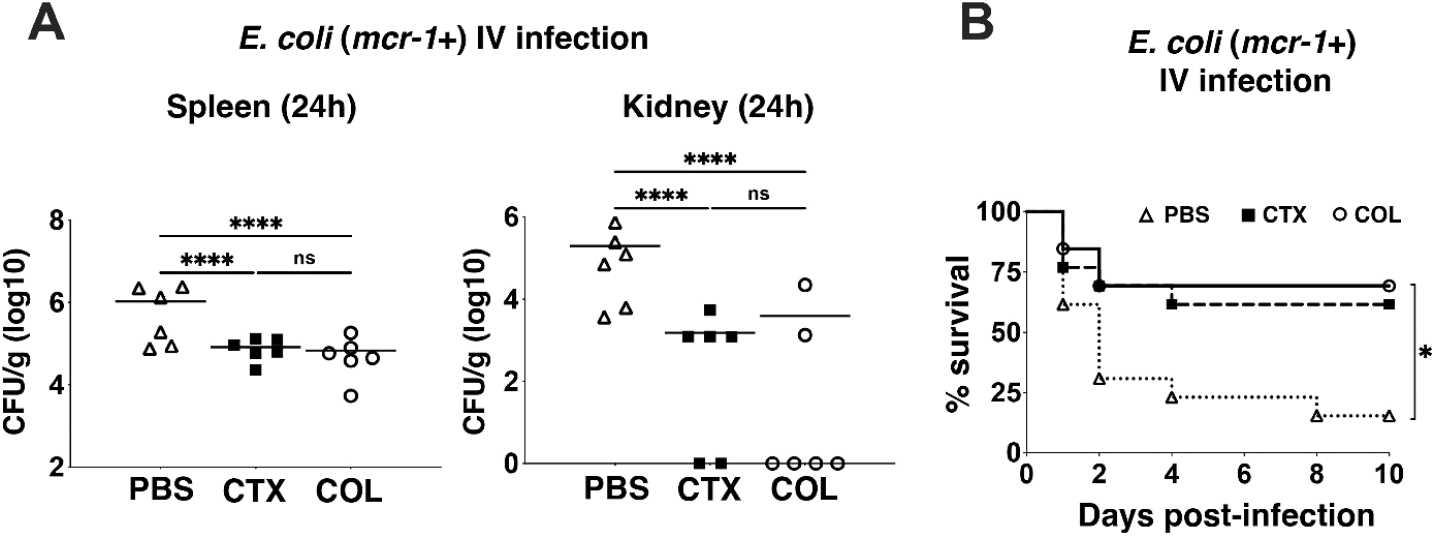
COL monotherapy reduces bacterial load and increases survival in a murine model of *mcr-1+ E. coli* bacteremia. (A) C57BL/6J mice were intravenously (iv) infected with 2 x 10^7^ CFU of *E. coli (mcr-1*) and treated subcutaneously with PBS (100 L, ⍰), CTX (50 mg/kg, ⍰) or COL (20 mg/kg, □) every 12 h for 2 doses (n = 6 per group). Bacterial loads recovered from spleen or kidney at 24 h plotted as CFU/g organ tissue. (B) Survival of C57BL/6J mice iv infected with 1 x 109 CFU of EC (*mcr*-1) and sc treated with a single dose of PBS (100 μL, ⍰), CTX (50 mg/kg, ⍰) or COL (20 mg/kg, ⍰) 1 h post infection (n = 10 per group). *P 0.05, **P ≤ 0.01 and ****P ≤ 0.0001; one-way ANOVA for CFU studies and log-rank test for survival.

## DISCUSSION

The experiments we present highlighting the unrecognized antibiotic action of COL against *mcr-1+* Gram-negative bacteria are straightforward, encompassing MIC testing, kinetic killing assays, synergy testing, and *in vivo* treatment in an animal model, yet their overarching medical implications are acute and significant. In the present era of expanding MDR and extreme drugresistant (XDR) Gram-negative bacterial infections, polymyxins (COL and PMB) have reemerged as a critical and last line antimicrobial therapy. The recent identification and spread of *mcr-1*, the plasmid-borne and horizontally transmissible COL resistance gene to superbugs resistant to nearly all antibiotics in our existing antimicrobial arsenal, has raised a global public health alarm. If a clinical strain isolate presents a COL MIC in AST testing above the published CLSI or EUCAST cutoffs for susceptibility, or if *mcr-1* is detected through FDA-approved PCR-based diagnostics such as the BioFire® or Verigene® blood culture panels (44–46), then clinicians, including infectious diseases specialists, undoubtedly refrain from using COL in the treatment regimen for the infected patient. This logic and calculus, however, is firmly rooted in information provided by AST performed in nonphysiologic bacterialogical growth media.

Despite lack of appreciable therapeutic antibacterial activity in the standard AST paradigm, our study revealed striking bactericidal activity of COL against several clinically important *mcr+* Gram-negative pathogens when testing was performed in the supplemented tissue culture medium, RPMI(10%LB). COL activity against *mcr-1+* EC required the crucial physiologic buffer bicarbonate, and paralleled results obtained *ex vivo* in human whole blood and *in vivo* in a murine model of bacteremia. Indeed, COL was non-inferior to the comparator CTX (identified to have a therapeutic MIC in CA-MHB) in murine survival and bacterial burden from harvested organs.

In the 1970s, decades before the discovery of *mcr-1*, a series of classical studies found additive or synergistic effects of PMB and human serum against Gram-negative pathogens including *E. coli, Serratia marcescens*, and *Salmonella typhimurium* whether or not the strain exhibited intrinsic (chromosomal) resistance to polymyxins [28–30]. We found COL strongly potentiated serum killing of mcr+ EC, KP and SE and that this effect was eliminated by heat-inactivation of complement, a finding corroborated by flow cytometry and fluorescence microscopy demonstration of increased serum complement binding to bacterial surface in the presence of COL. Serum typically constitutes 55% of human blood volume, and with just 10% serum supplementation we found marked sensitization of *mcr+* bacteria at COL concentrations (0.25 or 0.5 mg/L), well below the EUCAST clinical breakpoint for susceptibility (≤ 2 mg/L). Indeed addition of 10% serum allowed COL to kill *mcr+* strains in both physiological media RPMI(10%LB) and standard CA-MHB bacteriological media in our MIC assays (Tables 1 and 2), suggesting a simple modification to standard clinical microbiology laboratory workflows is at hand to discover these hidden and potentially clinically impactful antibiotic activities.

As familiar therapeutic alternatives dwindle to a precarious few for established or emerging bacterial pathogens, including several MDR Gram-species of urgent concern, it is incumbent upon clinical microbiologists, infectious diseases researchers and clinicians to ensure approved antibiotic agents are evaluated in the most comprehensive and holistic manner. Upon testing *mcr+* strains of Gram-negative pathogens EC, KP, and SE in physiological media and/or the presence of human serum, continued susceptibility to COL was apparent at drug levels readily attained with standard dosing, and validated by potent COL killing of each *mcr+* pathogen in freshly isolated human blood and *mcr+* EC in a murine bacteremia model. We hope these observations can inspire careful clinical trial design to determine if continued COL administration can contribute to successful therapeutic regimens in serious human *mcr+* Gram-negative bacterial infections, and that similar analyses can be applied to other drug and pathogen combinations to ensure familiar antibiotics are not being prematurely declared obsolete.

## METHODS

### Bacterial strains, media, and antibiotics

Twelve Enterobacteriaceae strains harboring *mcr-1* to *mcr-4* chosen from the Isolates with New or Novel Antibiotic Resistance Panel (CDC and FDA Antibiotic Resistance Isolate Bank) (50) were investigated for susceptibility to the cationic peptide antibiotics COL and PMB. Three *mcr-1+* strains, EC (AR Bank #0494), KP (AR Bank #0497), and SE (AR Bank #0496), were used in subsequent experiments. Isolates were stored in LB broth □+□ 50% glycerol at −80°C until use. COL, PMB and CTX were purchased from Sigma–Aldrich. Bacteriological medium Mueller–Hinton broth (Difco) was supplemented with 20–25 mg/L Ca^2+^ and 10–12.5 mg/L Mg^2+^ (CA-MHB). Tissue culture medium Roswell Park Memorial Institute 1640 (RPMI) (ThermoFisher Scientific) was supplemented with 10% Luria-Bertani broth (LB) (Hardy Diagnostics) yielding RPMI□ (10%LB).

### Antibiotic susceptibility assay

MIC assays of COL and PMB were performed per CLSI and EUCAST guidelines using the broth microdilution methodology with the recommended standard media (CA-MHB), the alternative cell culture media containing bicarbonate buffer (RPMI(10%LB)), and a final bacterial concentration of 5 x 10^5^ CFU/mL. Further assays were performed in the presence or absence of 10% human serum or 10% heat inactivated human serum. MICs were determined based on visual turbidity and absorbance (OD_600_) following 20 – 22 h incubation at 37°C.

### Kinetic killing assay

EC, KP and SE were grown overnight in LB broth, washed twice and diluted in CA-MHB, CA-MHB + COL (1x MIC), RPMI(10%LB), and RPMI(10%LB) + COL (1x MIC) to 5□×□10^5^ CFU/mL using a 96-well round bottom plate, in triplicate wells at a final volume of 100 μl/well, and incubated at 37°C with shaking for 24 h. Samples were collected at 0, 2, 4, 8 and 24 h of incubation, serially diluted in sterile PBS, and plated on LB agar for CFU enumeration. Bactericidal activity was defined as a reduction in viable bacteria by ≥3 log_10_ CFU/mL at 24 h compared to the starting inoculum.

### Bicarbonate supplementation assay

Overnight bacterial cultures (EC, KP, SE) grown in LB broth were washed twice and diluted to 5 x 10^5^ CFU/mL in CA-MHB (with 100 mM Tris to maintain pH ~7.3) or RPMI(10%LB) with 0, 10 or 25 mM of NaHCO_3_ (spanning physiologic NaHCO_3_ concentrations seen in humans) and 0, 1 or 2 μg/mL of COL in a total volume of 100 μL, in triplicate wells of a 96-well round bottom plate. Plates were incubated at 37°C shaking for 24 h prior to being serially diluted in sterile PBS, and plated on LB agar for CFU enumeration. Surviving bacteria were expressed as log_10_ CFU/mL.

### NPN (N-phenyl-1-naphthylamine) outer membrane permeabilization assay

Overnight cultures of EC grown in LB broth at 37°C in a shaking incubator were washed twice with PBS via centrifugation at 4000 rpm, and then resuspended in 10 mL of CA-MHB, CA-MHB + COL (1 μg/mL), RPMI(10%LB), or RPMI(10%LB) + COL (1 μg/mL) at an OD_600_ = 0.4. Cultures were incubated for 2 h at 37°C in a shaker, centrifuged at 4000 rpm for 5 min, and resuspended in 2 mL of 10 mM Tris buffer (pH 8.0). The concentrated 2 mL cultures were used to prepare 4 mL bacterial stocks in 10 mM Tris at OD_600_ = 0.4. Assays were performed at a final volume of 200 μL in triplicate, and using a 96 well flat bottom plate. The four conditions tested were: (1) Bacteria (100 μL) + 40 μM NPN (50 μL) + 10 mM Tris (50 μL); (2) Bacteria (100 μL) + 40 μM NPN (50 μL) + 10 mM EDTA (50 μL); (3) Bacteria (100 μL) + 10 mM EDTA (50 μL) + 10 mM Tris (50 μL); and (4) 10 mM EDTA (50 μL) + 40 μM NPN (50 μL) + 10 mM Tris (100 μL) Following addition of all components and mixing, fluorescence was immediately read at an excitation and emission of 250 nm and 420 nm respectively. To obtain the NPN fluorescence signal measured, conditions 3 and 4 (background) were subtracted from conditions 1 and 2 respectively. Following subtraction of background, the % NPN outer membrane permeabilization in the presence of EDTA was determined by dividing the measured NPN fluorescence signal of condition 1 (Bacteria + NPN) from condition 2 (Bacteria + NPN + EDTA) (24, 51).

### Cytochrome C binding assay

Bacterial surface charge was determined as previously described with the following modifications (52, 53). Briefly, stationary phase bacteria (EC) grown overnight in CA-MHB or RPMI(10%LB) were washed twice using 20 mM MOPS (pH 7.0), then resuspended to an OD_600_ = 0.7 in 500 μL of the same buffer with 0.5 mg/mL cytochrome c. The suspension was then incubated at room temperature in the dark for 10 min prior to being pelleted. Next the supernatant (in 100 μL aliquots) was transferred to a 96-well flat bottom plate, and the amount of unbound cytochrome c was measured spectrophotometrically at 530 nM. Percent bound cytochrome c was determined using a standard curve derived from a regression line and based on samples containing MOPS buffer and varying concentrations of cytochrome c (from 0.0009 to 0.5 mg/mL), and bacterial samples co-incubated with MOPS buffer only.

### Whole blood killing

Stationary phase bacteria (EC, KP and SE) were washed twice, diluted to an inoculum of 1 - 2 x 10^6^ CFU in 50 μL PBS and mixed with 400 μL heparinized human whole blood and 50 μL PBS with or without COL (at ¼ x MIC, ½ x MIC or 1 x MIC identified in supplemented RPMI) in siliconized tubes. Tubes were incubated at 37°C and rotated for 1 and 2 h. After incubation, the infected blood was serially diluted using sterile PBS and 0.025% Triton-X 100, and plated on LB agar plates. Percentage bacterial survival was defined as the number of CFU enumerated divided by the initial bacterial inoculum x 100%.

### Serum complement killing

Human serum was pooled from three healthy donors, stored as small aliquots at −80°C for no longer than 3 months, and thawed on the day of the experiment (kept at 4°C for ~1 h before use). Heat inactivated normal human serum (serum heated to 56°C for 30 min prior to use) served as a control. Overnight bacterial cultures (EC, KP, SE) grown in LB broth were washed twice and diluted to ~2 x 10^8^ CFU/mL in RPMI 1640. To measure bacterial survival, 1 - 2 x 10^6^ bacterial CFU in 20 μL were added to RPMI 1640 ± 10% pooled human serum, and varying concentrations of COL (0 mg/L, ¼ x MIC, ½ x MIC or 1 x MIC identified in RPMI) to a final volume of 200 μL in siliconized tubes rotated at 37°C. Samples were collected at 1 and 2 h, serially diluted in PBS, and plated on LB agar for CFU enumeration. Percentage survival was defined as the number of CFU enumerated divided by the initial bacterial inoculum x 100%.

### Complement deposition

C3 deposition on the bacterial surface was determined as previously described (54, 55) with minor modifications. Overnight bacterial cultures (EC, KP, SE) grown in LB broth were washed 3X with PBS and resuspended in RPMI 1640. Next, 1 - 2 x 10^6^ CFU of bacteria in 20 μL was added to RPMI 1640 containing no human serum, 20% human serum +/- 1 x MIC of COL or 20% heat inactivated human serum +/- 1 x MIC of COL to a final volume of 200 μL. Bacterial samples were then incubated at 37°C for 1 (EC) or 2 h (KP and SE). After incubation, bacteria were washed with PBS 3X, stained with a 1:200 dilution of LIVE/DEAD™ Fixable Violet Dead Cell Stain (Invitrogen) in PBS, and then incubated for 30 min at RT. Afterwards, bacteria were washed with PBS 3X and fixed with 4% paraformaldehyde for 20 min at RT. Bacteria were then washed 3X with PBS, incubated for 30 min at RT with blocking buffer (10% BSA + 0.01% of NaN_3_ in PBS), pelleted at 4000 RPM for 10 min, then resuspended in 1:500 dilution of rabbit anti-C3/C3b/C3c 1° Ab (Proteintech, 21337-1-Ab) with blocking buffer and incubated for 30 min at 4°C. Bacteria were then washed with PBS 3X, resuspended in 1:5000 dilution of Alexa Fluor 488 labeled goat anti-rabbit IgG 2° Ab (Invitrogen, A-11070) with blocking buffer and incubated for 30 min at 4°C. Lastly, bacteria were washed with PBS 3X before being resuspended in PBS and immediately analyzed using the BD FACS Canto II flow cytometer (BD Biosciences). Forward and side scatter were used to exclude debris and aggregates, and 10,000 gated events were recorded for each sample. Data were then analyzed with FlowJo v10.2 software (FlowJo, LLC) to identify the mean and median fluorescence intensity. Negative controls including bacteria without serum and bacteria stained with solely 2° Ab were used for setting gate boundaries. Additionally, heat killed bacteria were used as a positive control for the dead cell stain. Note: Each condition and controls were performed in triplicate, and all incubations were maintained in a light free environment.

### Immunofluorescence

For fluorescence microscopy, EC (*mcr-1*) were prepared and incubated at 37°C for 1 h as described above. Bacteria were then washed 3X with PBS and allowed to bind onto poly-D-Lysine-coated coverslips for 1 h (Corning). Next, coverslip bound bacteria were fixed with 4% paraformaldehyde in PBS, coverslips blocked by incubation in blocking buffer (5% BSA + 0.02% of NaN_3_ in PBS) for 30 min at RT, and stained with 1:500 dilution of rabbit anti-C3/C3b/C3c 1° Ab (Proteintech, 21337-1-Ab) for 1 h at RT. Coverslips were washed 3X in PBS, stained with 1:1000 Alexa Fluor 488 labeled goat anti-rabbit IgG 2° Ab (Invitrogen, A-11070) with blocking buffer and incubated for 1 h at RT before washing again 3X with PBS.

Coverslips were stained with 2 μM Hoescht dye (Thermo Fisher Scientific) and incubated for 15 min at RT, washed 3X with PBS and then mounted onto slides with ProLong Gold Antifade Mountant (Invitrogen). Images were acquired using a Leica SP8 Super Resolution Confocal microscope with 100X objective and fluorescence signals were quantified using ImageJ. The outline of 120 bacteria were traced, and the integrated density and cell area recorded. Mean fluorescence of 4 surrounding background regions were also measured. The corrected total cellular fluorescence (CTCF) was calculated as CTCF = integrated density – (area of selected cell x mean fluorescence of background regions). The relative fold CTCF was calculated and reported as fold fluorescence compared to untreated EC.

### Murine intravenous infection model

Mice were housed in a specific pathogen free (SPF) facility on a 12 h light/dark cycle in pre-bedded corn cob disposable cages (Innovive), fed a 2020X diet (Envigo), and received acidified water ad libitum. UCSD vivarium staff randomized mice 48-72 h prior to experimentation into cages with no more than 5 mice per cage. The *in vivo* activity of COL and CTX against *mcr-1+* EC harboring *mcr-1* was evaluated using an immunocompetent murine intravenous challenge (56). Eight- to 10-week-old female C57Bl/6J mice (Jackson Labs) were infected IV via the right retro-orbital venous sinus with 2 x 10^7^ CFU of EC in 100 μL PBS for the bacterial burden study, and treated with COL (20 mg/kg, subcutaneously (sc), n = 6), CTX (50 mg/kg, sc, n = 6) or PBS (100 μL, sc, n = 6) every 12 hours (at 1 and 13 h post infection) for a total of 2 doses. Mice were euthanized 24 h post infection by CO_2_ asphyxiation and cervical dislocation. Organs (right kidney and spleen) were aseptically collected, weighed, homogenized, serially diluted and plated on LB agar for CFU enumeration following 24 h of incubation at 37°C. To assess survival, all operations of infection were similar, except mice were infected intravenously with 1 x 10^9^ CFU of EC (*mcr-1*) in 100 μL PBS and treated with only one dose of COL (20 mg/kg, sc, n = 13), CTX (50 mg/kg, sc, n = 13) or PBS (100 μL, sc, n = 13) 1 h post infection. Survival was monitored every 12 h till the endpoint of the experiment (defined as 10 days post infection), and date and time of death was recorded for each mouse.

### Statistics

Statistical analyses were conducted using Prism 9.0 (GraphPad Software Inc.) on at least three independent experiments unless otherwise stated. Data are represented as the mean ± standard error of the mean (SEM) where applicable and *P* values < 0.05 were regarded to be statistically significant. Sample size and information about statistical tests are reported in the Methods section and figure legends.

### Study approval

Human blood and serum were obtained from healthy donors with informed consent under a simple phlebotomy IRB protocol #131002 approved by the UC San Diego Human Research Protection Program. All murine infection and antibiotic treatment studies were conducted in compliance with federal Animal Welfare Act regulations and the UC San Diego Institutional Animal Care and Use Committee (IACUC) under approved Protocol #S00227M.

## Author contributions

**MK** contributed to research study design, experiments (MIC, checkerboards, kinetic killing, HCO3- sensitization, membrane permeabilization, serum and whole blood killing, complement studies, microscopy, mouse infections), data acquisition and analysis, figure generation, and manuscript writing. **AR** contributed to research study design and conducted experiments (complement binding, flow cytometry, fluorescence microscopy), data acquisition and analysis, figure generation, and manuscript writing. **AF** conducted experiments (kinetic killing, HCO3- sensitization, membrane permeabilization), data acquisition and analysis. **SU** conducted murine models of infection, data acquisition and analysis, and figure generation, **GB**, **VNil**, and **MC** conducted preliminary experiments (MIC, whole blood killing), data acquisition and analysis. **SD**, **HS**, and **GS** contributed to research study design data acquisition and analysis. **VN** contributed to the overall research study design, data, analysis, figure generation, and to the writing and editing of the manuscript.

## Acknowledgments

This work was supported by the National Institute of Health grants U54-HD090259 (to GS and VN), U01-AI124316 (to MK, GS, and VN), KL2-TR001444 (to MK), and R01AI145310 (to MK and VN). AF was supported by the NIH-funded San Diego IRACDA K12 grant program (GM068524).

